# A fitness cost resulting from *Hamiltonella defensa* infection is associated with altered probing and feeding behaviour in *Rhopalosiphum padi*

**DOI:** 10.1101/652974

**Authors:** Daniel. J. Leybourne, Tracy. A. Valentine, Jorunn. I. B. Bos, Alison. J. Karley

## Abstract

Aphids frequently associate with facultative endosymbiotic bacteria which influence aphid physiology in myriad ways. Endosymbiont infection can increase aphid resistance against parasitoids and pathogens, modulate plant responses to aphid feeding, and promote aphid virulence. These endosymbiotic relationships can also decrease aphid fitness in the absence of natural enemies or when feeding on less suitable plant types. Here, we use the Electrical Penetration Graph (EPG) technique to monitor feeding behaviour of four genetically-similar clonal lines of a cereal-feeding aphid, *Rhopalosiphum padi*, differentially infected (+/−) with the facultative protective endosymbiont, *Hamiltonella defensa*, to understand how physiological processes at the aphid-plant interface are affected by endosymbiont infection. Endosymbiont-infected aphids exhibited altered probing and feeding patterns compared with uninfected aphids, characterised by a two-fold increase in the number of plant cell punctures, a 50% reduction in the duration of each cellular puncture, and a greater probability of achieving sustained ingestion of plant phloem. Feeding behaviour was altered further by host plant identity: endosymbiont-infected aphids spent less time probing into plant tissue, required twice as many probes into plant tissue to reach plant phloem, and showed a 44% reduction in phloem ingestion when feeding on the partially-resistant wild relative of barley, *Hordeum spontaneum* 5, compared with a commercial barley cultivar. These observations might explain reduced growth of *H. defensa*-infected aphids on the former host plant. This study is the first to demonstrate a physiological mechanism at the aphid-plant interface contributing to endosymbiont effects on aphid fitness on different quality plants through altered aphid feeding behaviour.

**Summary:** Reduced performance of aphids infected with a common facultative endosymbiont on poor quality plants may be explained by changes in aphid probing behaviour and decreased phloem sap ingestion.

## Introduction

Aphids are major pests of agricultural and horticultural crops with worldwide distribution (Van Emden and Harrington, 2017). To feed they probe into plants using specialised mouthparts known as stylets, with the aim of establishing a feeding site in the plant phloem. While probing into plant tissues, aphids can transmit plant viruses (Powell, 2005), which are the main cause of economic crop losses resulting from aphid infestation. Most aphid species harbour the obligate bacterial endosymbiont *Buchnera aphidicola*, which supplies aphids with essential amino acids they are unable to biosynthesise (Sasaki et al., 1991; Douglas and Prosser, 1992). Additional co-obligatory symbiotic relationships have been described with other endosymbiont species, including *Wolbachia sp.* in the banana aphid, *Pentalonia nigronervosa* (De Clerck et al., 2015)(although see Manzano-Marín, 2019 preprint for updated analysis), and with *Serratia symbiotica* in multiple species of the *Cinara* genus (Meseguer et al., 2017). In most other aphid species, however, these co-infecting endosymbionts are not essential for survival. Alongside obligatory endosymbiotic relationships, aphids can form facultative (non-essential) endosymbiotic relationships with a range of microorganisms.

The diversity and frequency of infection with facultative endosymbionts can vary considerably between and within aphid species (de la Peña et al., 2014; Henry et al., 2015; Zytynska and Weisser, 2016; Guo et al., 2019). The eleven most common facultative endosymbionts are *Hamiltonella defensa, Regiella insecticola, S. symbiotica, Rickettsia sp., Ricketsiella sp*, the *Pea Aphid X-type Symbiont (PAXS), Spiroplasma sp., Wolbachia sp., Arsenophonus sp., Sitobion miscanthis L-type Symbiont (SMLS)*, and *Orientia-Like Organism (OLO)* (Oliver et al., 2006; Castañeda et al., 2010; Tsuchida et al., 2010; Łukasik et al., 2013a; de la Peña et al., 2014; Leybourne et al., 2018). A meta-analysis of endosymbiont infection frequencies by Zytynska and Weisser (2016) assessed the proportion of aphid species shown to harbour *H. defensa, R. insecticola, S. symbiotica, Rickettsia sp., Spiroplasma sp., PAXS, Arsenophonus sp.* and *Wolbachia sp.* infections. The number of aphid species tested for each endosymbiont varied from 14 aphid species tested for PAXS infection, to 190 aphid species tested for *Wolbachia* infections (Zytynska and Weisser, 2016). This analysis found that the most frequently detected endosymbiont in aphids is *Serratia symbiotica* (47% of the 156 aphid species tested were infected) with *Arsenophonus sp.* the least frequently detected (7% of the 131 aphid species tested were infected) (Zytynska and Weisser, 2016).

The benefits of aphid infection with nine of these endosymbionts has recently been reviewed by Guo et al. (2017) and a meta-analysis of the costs and benefits of facultative endosymbiont infection has recently been conducted by Zytynska et al. (2019 preprint). Beneficial traits conferred to the aphid by the endosymbionts include protection against parasitism by Braconid wasps (*H. defensa* and *R. insecticola;* Hansen et al. (2012); Leybourne et al. (2018)), protection against entomopathogenic fungi (*R. insecticola, Rickettsia sp., Ricketsiella sp.* and *Spiroplasma sp*.; Łukasik et al. (2013b)), host-plant adaptation (*Arsenophonus sp.;* Wagner et al. (2015)), heat tolerance (*S*. *symbiotica* and *H. defensa*, alongside *B. aphidicola* mutations; Russell and Moran (2006); Dunbar et al. (2007)), morphological changes in insect colour (*Ricketsiella sp.;* Tsuchida et al. (2010); Nikoh et al. (2018)), and enhanced aphid virulence (mixed symbiont communities; Luna et al. (2018)). Infection with endosymbionts can, however, result in negative fitness consequences for the aphid host, including decreased growth (*Rickettsia sp.;* Sakurai et al., 2005), reduced fecundity (*Spiroplasma sp., H. defensa*, and *S*. *symbiotica*; Chen et al. (2000); Castañeda et al. (2010); Li et al. (2018); Mathé-Hubert et al. (2019)), shorter aphid longevity (*Spiroplasma sp.* and *S*. *symbiotica*; Chen et al. (2000); Mathé-Hubert et al. (2019)), lower adult mass (*S*. *symbiotica;* Skaljac et al. (2018)), and increased susceptibility to insecticide (*S*. *symbiotica;* Skaljac et al. (2018)).

Around one-third of 154 aphid species assessed for endosymbiont presence have been reported to harbour *H. defensa* (Zytynska and Weisser, 2016). Amongst cereal-feeding aphids, the proportion of the bird-cherry oat aphid, *Rhopalosiphum padi*, populations infected with the defensive endosymbiont, *H. defensa*, is around 10.8% (63/585 individuals; Guo et al., (2019)). The primary trait conferred to aphids infected with *H. defensa* is protection against parasitism by Braconid wasps (Oliver and Higashi, 2019). This protective phenotype often depends on the association of *H. defensa* with virulent strains of the bacteriophage *APSE (Acyrthosiphon pisum Secondary Endosymbiont)* which secretes toxins that arrest the development of the parasitoid wasp larvae (Brandt et al., 2017). A recent study has shown that *H. defensa* infection can alter interactions at the aphid-plant interface by influencing aphid probing behaviour (Angelella et al., 2018). This observation suggests that altered aphid probing behaviour could affect aphid fitness by altering feeding success and thus potentially impacting on aphid nutrition. Indeed, *H. defensa*-infection has been shown to have consequences for aphid fitness (Castañeda et al., 2010; Vorburger and Gouskov, 2011; Li et al., 2018; Zytynska et al., 2019 preprint), and examining these symbiont effects in relation to aphid probing behaviour could elucidate the mechanistic processes contributing towards symbiont-associated fitness consequences.

The Electrical Penetration Graph (EPG) technique is an electrophysiological tool used to monitor the probing and feeding behaviour of sap-feeding insects (Tjallingii, 1985; Tjallingii and Esch, 1993; Prado and Tjallingii, 1994) and has been used successfully to monitor the feeding and probing behaviour of aphids (Greenslade et al., 2016), whiteflies (Chesnais and Mauck, 2018), psyllids (Civolani et al., 2011) and planthoppers (He et al., 2011). The technique is based on an electrical circuit which is made by inserting conductive copper probes into the soil around the plant and adhering conductive wire onto the dorsum of the aphids (Tjallingii, 1978; Tjallingii, 1985). Both probes are connected to a data logger and computational software. An electrical current is passed through the circuit, which is closed when the aphid stylet comes into contact with plant tissue, and the resulting electrical waveforms can be characterised to provide information on aphid stylet activities (probing and feeding behaviour) (Kimmins and Tjallingii, 1985; Tjallingii and Esch, 1993; Prado and Tjallingii, 1994; Tjallingii et al., 2010). Different electrical waveforms obtained from EPG recordings in aphids correspond with stylet activities in specific cell layers (Sarria et al., 2009), including in the mesophyll tissue (pathway, C, phase), xylem (G phase - xylem ingestion), and phloem (E phase, split into E1 - salivation into the phloem, E2 - ingestion of phloem sap, and sE2 - prolonged ingestion of phloem for a period of time > 10 mins). Alongside these main phases, waveforms can also relate to difficulties in stylet penetration of plant tissue (F phase), salivation into the extracellular space (E1e), and stylet puncture of cells within the mesophyll tissue (pd) (Sarria et al., 2009; Tjallingii et al., 2010). A primary use of the EPG technique has been to identify plant tissue types involved in plant resistance against sap-feeding pests (Alvarez et al., 2006; Greenslade et al., 2016). However, the EPG technique can also be employed to examine insect physiological responses to a myriad of biotic and abiotic factors, such as environmental stress (Ponder et al., 2000), plant disease status (Angelella et al., 2018), plant association with mycorrhiza (Simon et al., 2017), and disruption of the obligate aphid endosymbiont *B. aphidicola* (Machado-Assefh and Alvarez, 2018).

In the current study we use the EPG technique to examine aphid feeding on a susceptible modern cultivar of barley, *Hordeum vulgare* cv. Concerto, and a wild relative of barley with partial-resistance against aphids, *H. spontaneum* 5 (Hsp5) (Delp et al., 2009). Hsp5 is particularly unsuitable as a host for *H. defensa*-infected *R. padi* (Leybourne et al., 2018). We analysed aphid feeding behaviour to test two complementary hypotheses: 1) that infection with *H. defensa* can lead to altered aphid probing and feeding behaviour; 2) that differential aphid probing and feeding behaviour between uninfected and *H. defensa*-infected aphids is a key contributor towards the decreased fitness of *H. defensa*- infected aphids feeding on partially-resistant Hsp5. In support of the first hypothesis we observed that, irrespective of plant type, aphids infected with *H. defensa* exhibited differential physiological behaviour relating to stylet interactions with the mesophyll tissue and the phloem sap. For example, infected aphids probed intracellularly within mesophyll tissue more frequently, salivated into the phloem for a shorter period of time, and were more likely to establish sustained phloem feeding. Furthermore, in support of the second hypothesis, we also observed that, compared with uninfected aphids, *H. defensa*-infected aphids on Hsp5 spent less time probing into plant tissue, required twice as many tissue probes in order to reach the phloem sap, and showed a significant reduction in time spent ingesting phloem. Our work provides novel information about the mechanisms (decreased probing into plant tissue and reduced phloem access) that contribute towards the fitness consequences for *H. defensa*-infected aphids feeding on a less suitable plant host.

## Materials and Methods

### Plant growth and aphid rearing conditions

Barley seeds, *Hordeum vulgare* cv. Concerto (Linnaeus) (Concerto), and wild barley seeds, *H. spontaneum* (Linnaeus) 5 (Hsp5) were surface sterilised by washing in 2% (v/v) hypochlorite solution and rinsed in deionised water. Concerto seeds were kept in the dark at room temperature for 48 h to germinate whereas Hsp5 seeds were incubated at 4°C in the dark for 14 days to synchronise germination. Plants were grown to the true-leaf stage of development (1.2 on the scale described in Zadoks et al. (1974)) before use in aphid fitness and EPG experiments.

Asexual laboratory clonal cultures of the bird cherry-oat aphid, *Rhopalosiphum padi* (Linnaeus), were reared on one week old barley seedlings (cv. Optic) contained in ventilated cups and maintained at 18 ± 2°C and 16h:8h (daymight). *R. padi* lines were previously genotyped and characterised for the presence of facultative endosymbionts (Leybourne et al., 2018). Prior to experimentation, the presence of the defensive endosymbiont *Hamiltonella defensa* (Moran et al., 2005) was confirmed by PCR on a ProFlex PCR system (Applied Biosystems, UK) with PCR conditions as follows: 95°C for 2 min followed by 35 cycles of 95°C for 30s, 55°C for 30s and 72 °C for 3 min with a final extension stage of 72°C for 5 min; the final reaction solution consisted of 1X Green GoTaq^®^ reaction buffer (Promega, UK) containing 1 μmol forward primer (PABSF: 5’ - AGCGCAGTTTACTGAGTTCA - 3’ (Darby and Douglas, 2003), 1 μmol reverse primer (16SB1 5’ - TACGGYTACCTTGTTACGACTT -3’ (Fukatsu et al., 2000), 1.25 U GoTaq® DNA Polymerase (Promega, UK) and 1.5 mmol MgCI_2_.

### Electrical Penetration Graph (EPG) aphid feeding assessment

The DC-EPG technique (Tjallingii, 1978; Tjallingii, 1988) was employed to monitor probing and feeding behaviour of four *R. padi* lines representing one aphid genotype with differential infection with the aphid endosymbiont, *H. defensa:* DL 16/04 (*Hd+*), DL 16/05 (*Hd+*), DL 16/06 (*Hd*−) and DL 16/13 (*Hd*−). Recordings were taken over a six-hour period using a Giga-4 DC-EPG device (EPG Systems, The Netherlands). Aphids were adhered to aphid probes (a copper wire, 3 cm × 0.2 cm, soldered to a brass pin, tip width 0.2 cm) by attaching 3 cm of gold wire (20 μm diameter; EPG Systems, The Netherlands) to the aphid probe using water-based silver glue (EPG Systems, The Netherlands) and adhering the free end of the wire to the aphid dorsum using the same water-based adhesive. A plant probe (copper rod approximately 5 cm × 0.5 cm) was created by soldering the copper rod to electrical wire extending from the plant voltage output of the Giga-4 device. The wired aphid was attached to the Giga-4 device by placing the end of the brass pin into the EPG probe and the copper rod was then placed into the plant soil. Recordings were taken with a 1G Ω input resistance and a 50 × gain (Tjallingii, 1988), for six hours per read. The order of *R. padi* - plant combinations and allocation to EPG probe was randomised, and Stylet+D software (EPG Systems, The Netherlands) was used for data acquisition. Aphids were lowered onto plant leaves immediately after the recording started. All EPG recordings were taken in a grounded Faraday cage. Ten replicates were recorded for *H. defensa*-infected aphids on Concerto, 11 for *H. defensa*-infected aphids on Hsp5,14 for uninfected aphids on Concerto, and 14 for uninfected aphids on Hsp5

EPG waveforms were annotated using Stylet+A software (EPG Systems, The Netherlands) by assigning waveforms to np (non-probing), C (stylet penetration/pathway), pd (potential-drop/intercellular punctures), the pd sub-phases (pd-II1, pd-II2, pd-II3), E1e (extracellular saliva secretion), E1 (saliva secretion into phloem), E2 (saliva secretion and passive phloem ingestion), F (penetration difficulty) or G (xylem ingestion) phases (Tjallingii, 1988; Alvarez et al., 2006). Annotated waveforms were converted into time-series data using the excel workbook for automatic parameter calculation of EPG data (Sarria et al., 2009).

### Aphid fitness experiments

The aphid fitness study was split into two temporal blocks with seven fully-randomised sub-blocks within each temporal block; each sub-block consisted of one replicate for each plant-aphid-combination: two plant types (Hsp5, Concerto) x four aphid treatments (DL 16/04, DL 16/05, DL 16/06, DL 16/13) giving eight treatments with 14 replicates each. One apterous *R. padi* adult from the four aphid lines described above was taken from culture, placed in a perspex clip-cage (MacGillivray and Anderson, 1957), attached to the first true leaf and left to reproduce overnight. After 24 h the adult was removed and the resulting progeny were retained on the plant leaf; the mean mass of two nymphs was recorded at 48 h and 114 h and used to calculate the nymph mass gain over this 96 h period. We have previously characterised *R. padi* fitness in relation to *H. defensa*- infection (nymph mass, fecundity, survival) and detected a fitness consequence for nymph mass gain in *H. defensa*-infected aphids (Leybourne et al., 2018). As such, we only measured nymph mass in this current study.

### Statistical analysis

All statistical analyses were carried out using R Studio Desktop version 1.0.143 running R version 3.4.3 (R Core Team, 2017), with additional packages car v.2.1-4 (Fox and Weisberg, 2011), ggplot2 v.2.2.1 (Wickham, 2009), ggpubr v. 0.1.2 (Kassambara, 2017), lme4 v.1.1-13 (Bates et al., 2015), lmerTest v.2.0-33 (Kuznetsova et al., 2017), lsmeans v.2.27-62 (Lenth, 2016), multcomp v.1.4-8 (Hothorn et al., 2008), pastecs v.1.3.21 (Grosjean and Ibanez, 2014), and vegan v.2.4-6 (Oksanen et al., 2013).

Data were combined into two endosymbiont treatments: *H. defensa*-infected (comprising the DL 16/04 and DL 16/05 clonal lines) and *H. defensa*-uninfected (comprising the DL 16/06 and DL 16/13 clonal lines). Aphid feeding behaviour was first assessed globally by fitting a permutated multiple analysis of variance to the dataset. Individual feeding parameters from the EPG experiment and aphid juvenile mass gain from the aphid fitness experiment were then analysed in individual linear mixed effects models. Within each model, aphid clonal line was included as a nested factor within endosymbiont infection status to account for the use of multiple clonal lines of genetically similar aphids with differential endosymbiont infection status, as done previously (Leybourne et al., 2018). For the individual EPG parameters, EPG run (blocking factor) and the EPG probe used were included as random factors (there were 3 EPG probes used over the lifetime of the experiment). For the juvenile mass gain model, experimental block and temporal block were incorporated as random factors. All data were modelled against host plant, aphid endosymbiont infection status, and the interaction. χ^2^ analysis of deviance tests were used to analyse the final models for the individual EPG parameters and analysis of variance with type III Satterthwaite approximation for degrees of freedom was used to analyse the final aphid fitness model. Calculation of the Least Squares Means was used as a *post-hoc* test on all models with a significant interaction term. All final models were checked for model suitability by observing the fitted-residual plots.

## Results

We used the EPG technique to monitor the probing and feeding behaviour of genetically-similar *Rhopalosiphum padi* clonal lines with and without *H. defensa* infection when feeding on two host plants: susceptible barley (Concerto) and the partially-resistant wild relative (Hsp5). We tested two complementary hypotheses: 1) that infection with *H. defensa* can lead to altered aphid probing and feeding behaviour; 2) that differential aphid probing and feeding behaviour between uninfected and *H. defensa*-infected aphids is a key contributor towards the decreased fitness of *H. defensa*-infected aphids when feeding on partially-resistant Hsp5. We obtained 72 individual feeding parameters from the EPG analysis (displayed in tables 1 and 2, and supplementary tables 1 and 2). Global analysis of aphid feeding patterns indicated that aphid feeding behaviour was significantly affected by plant identity (F_1,43_= 3.19; p = 0.017) and the interaction between endosymbiont presence and plant identity (F_1,43_ = 2.71; p = 0.037). From the 72 parameters obtained, seven parameters were affected by plant identity alone (Table S1), however these parameters are not presented or discussed in detail here as we recently characterised *R. padi* feeding responses to plant identity in a separate study (see Leybourne et al. 2019). In support of hypothesis 1, a total of 11 parameters were primarily influenced by endosymbiont infection, and these were mainly associated with stylet intracellular punctures and interaction with the phloem (Table 1; Fig. 1). A further 15 parameters were differentially affected by the endosymbiont infection x host plant interaction (supporting hypothesis 2), and these involved stylet interactions with the plant surface, the mesophyll tissue, and the phloem (Table 2; Fig 2). The remaining 39 non-significant parameters are displayed in Table S2.

**Table 1:** Aphid feeding parameters (mean value ± standard error) that were significantly affected by the absence (-) or presence (-) of *Hamiltonella defensa* infection. Numbers in parenthesis indicate the total number of individuals which displayed each parameter, and the total number of individuals tested is indicated at the top of the column. Data marked with † display median alongside the upper and lower interquartile ranges.

| Description of Response Variable Assessed (transformation used) | Absence (-) or presence (+) of *Hamiltonella defensa* infection |  | Statistical results for each Explanatory Variable (generalised least square estimation models) |  |  |  |  |  |
| --- | --- | --- | --- | --- | --- | --- | --- | --- |
|  | *Hd* -ve (max = 28) | *Hd* +ve (max = 21) | Plant |  | Endosymbiont |  | Plant x Endosymbiont |  |
| Mean duration of each C (mesophyll pathway) phase (sqrt) | 860.14 ± 94.77 s (28) | 620.02 s ± 70.71 s (21) | $\chi^2_1 = 0.25$ | p = 0.611 | $\chi^2_1 = 4.16$ | p = 0.041 | $\chi^2_1 = 0.97$ | p = 0.323 |
| Total number of pd: potential drops (intracellular punctures) (sqrt) | 72.85 ± 6.83 (28) | 163.57 ± 26.02 (21) | $\chi^2_1 = 0.52$ | p = 0.470 | $\chi^2_1 = 18.49$ | p = <0.001 | $\chi^2_1 = 3.78$ | p = 0.051 |
| Mean duration of each pd (not transformed) | 4.53 s ± 0.17 s (28) | 2.66 s ± 0.28 s (21) | $\chi^2_1 = 0.03$ | p = 0.843 | $\chi^2_1 = 40.99$ | p = <0.001 | $\chi^2_1 = 0.74$ | p = 0.389 |
| Time from start of aphid probe into plant tissue to first pd (not transformed) | 529.41 ± 206.43 (28) | 117.75 s ± 36.49 s (21) | $\chi^2_1 = 2.09$ | p = 0.147 | $\chi^2_1 = 2.87$ | p = 0.008 | $\chi^2_1 = 1.58$ | p = 0.208 |
| Total number of pd in first hour (sqrt) | 15.35 ± 2.29 (28) | 61.52 ± 14.58 (21) | $\chi^2_1 = 0.02$ | p = 0.871 | $\chi^2_1 = 14.67$ | p = 0.001 | $\chi^2_1 = 1.25$ | p = 0.262 |
| Mean duration of each pd in first hour (not transformed) | 4.19 s ± 0.29 s (28) | 2.39 s ± 0.32 s (21) | $\chi^2_1 = 0.01$ | p = 0.977 | $\chi^2_1 = 16.62$ | p = <0.001 | $\chi^2_1 = 0.001$ | p = 0.994 |
| Mean duration of each pd in second hour (not transformed) | 4.81 ± 0.30 s (21) | 3.14 ± 0.47 (17) | $\chi^2_1 = 0.59$ | p = 0.439 | $\chi^2_1 = 10.01$ | p = 0.001 | $\chi^2_1 = 1.62$ | p = 0.202 |
| Mean duration of each pd in sixth hour (not transformed) | 4.84 ± 0.19 (13) | 3.57 ± 0.29 (13) | $\chi^2_1 = 0.92$ | p = 0.335 | $\chi^2_1 = 5.13$ | p = 0.023 | $\chi^2_1 = 4.32$ | p = 0.057 |
| Time spent in E1 (salivation into phloem) as a percentage of the total time spent in all phloem phases (not transformed) † | 16.37%<br>LQR: 6.90%<br>UQR: 45.61% (26) | 8.93%<br>LQR: 4.63%<br>UQR: 22.12% (20) | $\chi^2_1 = 1.13$ | p = 0.285 | $\chi^2_1 = 18.20$ | p = <0.001 | $\chi^2_1 = 2.02$ | p = 0.154 |
| E2 (phloem ingestion) index (not transformed) † | 29.44 %<br>LQR: 5.25%<br>UQR: 76.85% (24) | 41.36%<br>LQR: 8.95%<br>UQR: 78.67% (20) | $\chi^2_1 = 0.01$ | p = 0.987 | $\chi^2_1 = 6.72$ | p = 0.009 | $\chi^2_1 = 0.94$ | p = 0.329 |
| % of E2 phases which contained a period of sustained (>10min) phloem ingestion (not transformed) † | 45.00%<br>LQR: 27.08%<br>UQR: 100.00% (20) | 100.00%<br>LQR: 50.00%<br>UQR: 100.00% (17) | $\chi^2_1 = 0.37$ | p = 0.541 | $\chi^2_1 = 3.21$ | p = 0.047 | $\chi^2_1 = 0.90$ | p = 0.341 |

**Fig. 1:**
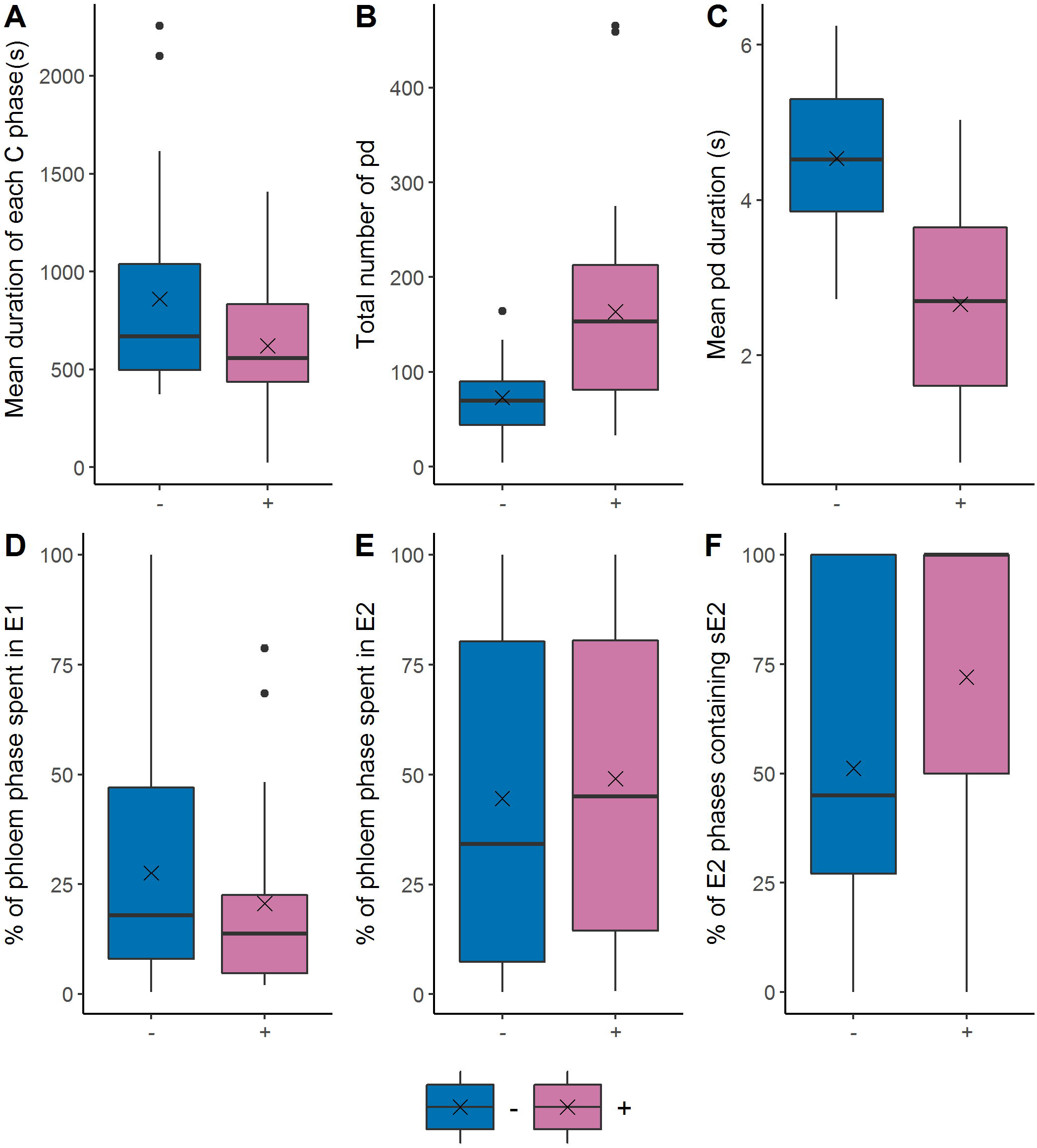
Aphid feeding parameters in the absence (−) and presence (+) of *Hamiltonella defensa*. - shows the combined data for both *H. defensa*-uninfected lines (DL 16/06; DL 16/13) and + shows the combined data for both *H. defensa*-infected lines (DL 16/04; DL 16/05). A-C: parameters associated with stylet puncturing of plant cells (intracellular punctures); D-F: parameters associated with stylet interaction with phloem sap. The black cross (“x”) on each plot shows the mean value.

**Table 2:**
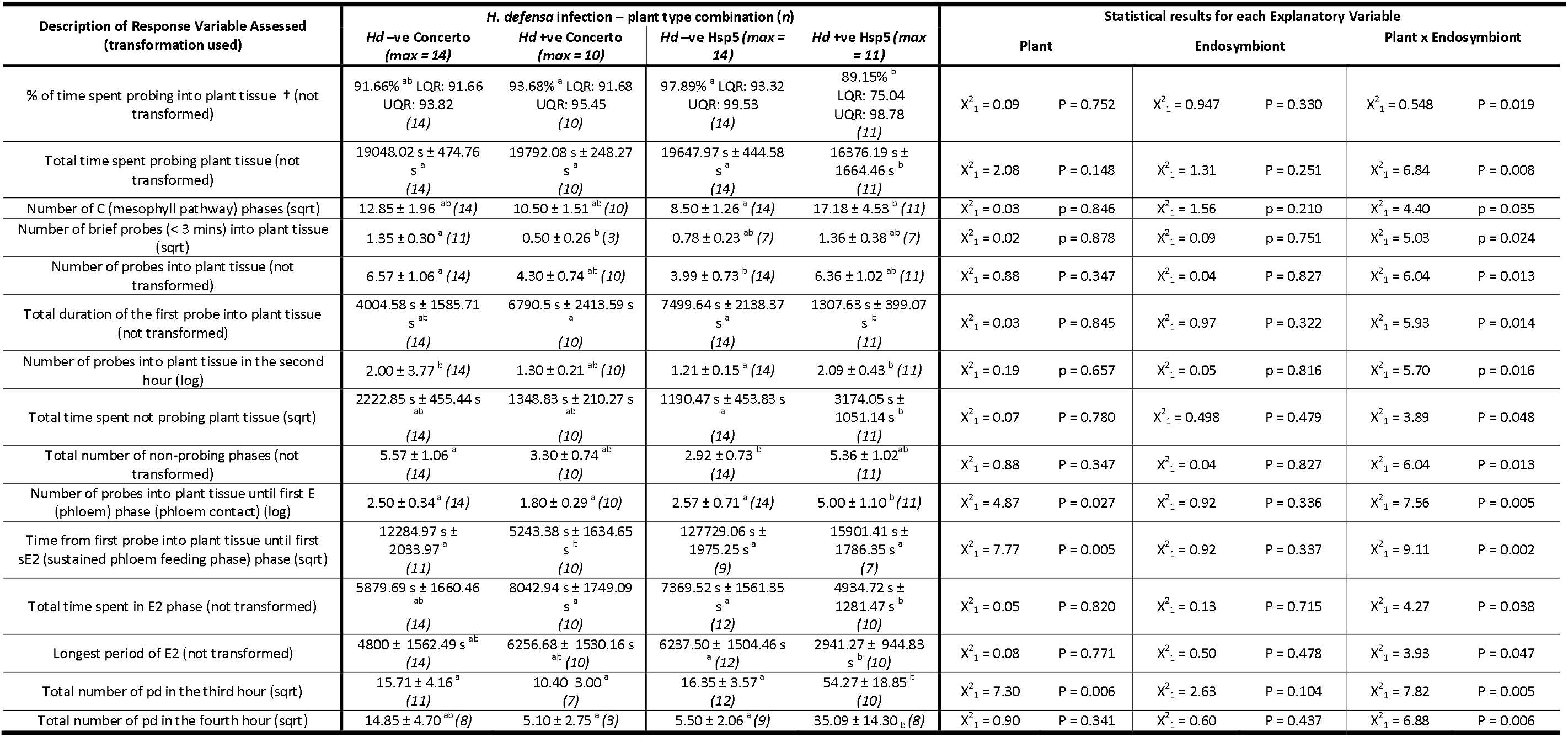
Aphid feeding parameters (mean value ± standard error) that were differentially affected by plant type and *Hamiltonella defensa* infection. Letters indicate which groups are significantly different based on pairwise comparisons using differences in the least square means analysis. Numbers in parenthesis indicate the total number of individuals which displayed each parameter, and the total number of individuals tested is indicated at the top of the column. Data marked with † display median alongside the upper and lower inter quartile ranges.

**Fig 2:**
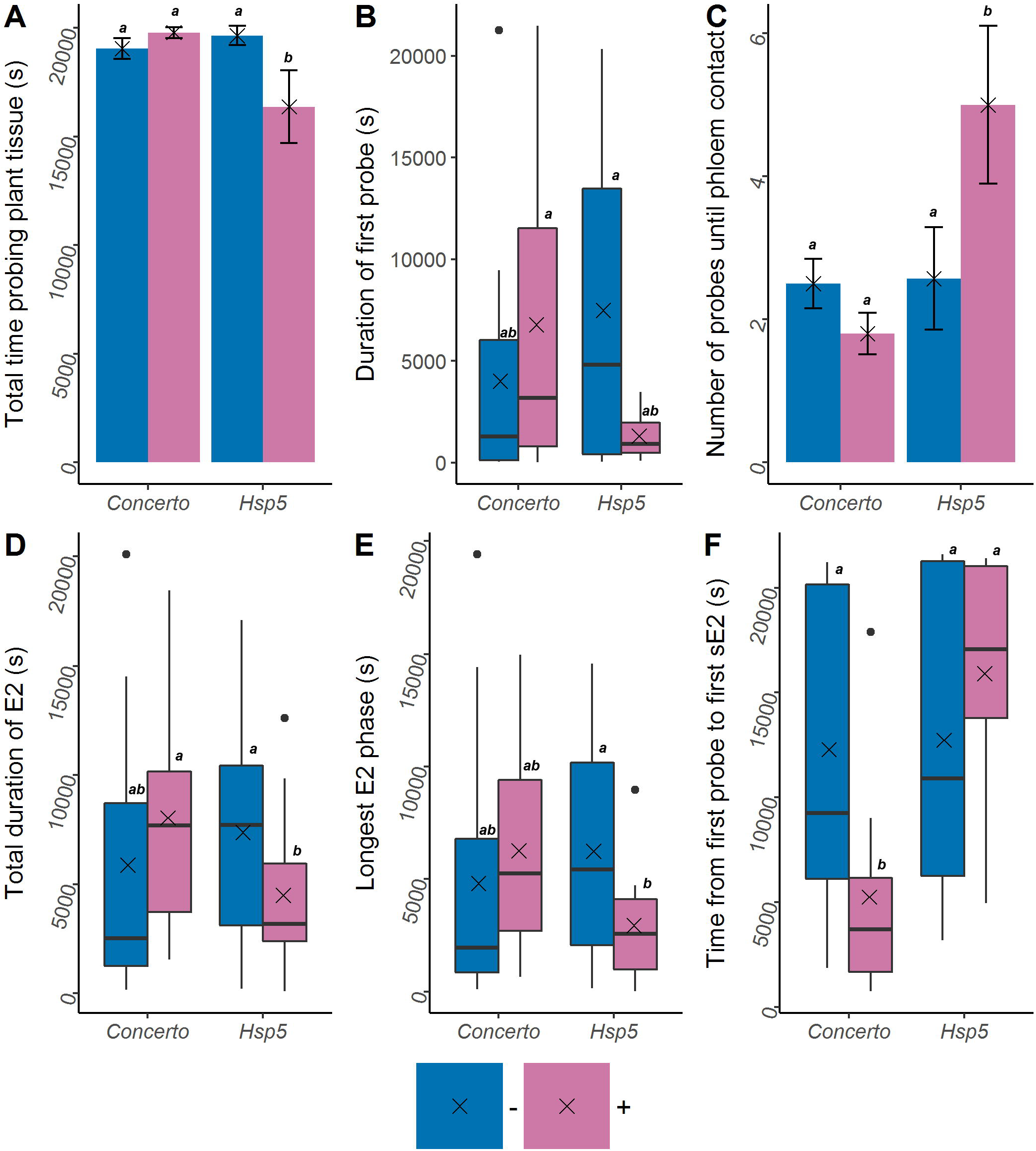
Aphid feeding parameters that were differentially affected by the absence (−) and presence (+) of *Hamiltonella defensa* infection on two plant hosts (susceptible modern barley cv. Concerto and the wild relative Hsp5). - shows the combined data for both *H. defensa*-uninfected lines (DL 16/06; DL 16/13) and + shows the combined data for both *H. defensa*-infected lines (DL 16/04; DL 16/05). Letters indicate which groups are significantly different based on pairwise comparisons using general linear hypotheses testing with single-step p-value adjustment. The black cross (“x”) on each plot shows the mean value.

### Cellular punctures and phloem feeding are more frequent in aphids infected with the defensive facultative endosymbiont Hamiltonella defensa

During the six-hour EPG recording, 11 feeding parameters were affected similarly by endosymbiont infection for aphids feeding on Concerto and Hsp5 (Table 1), supporting our first hypothesis that infection with *H. defensa* can lead to altered aphid probing and feeding behaviour. Most of these feeding parameters related to aphid stylet activities in the mesophyll tissue, specifically the frequency and duration of the exploratory intracellular punctures (EPG waveform pd) performed by aphids while probing into plant tissue, and stylet interaction with phloem sap. The average duration of each C phase (stylet interaction with and movement through the mesophyll tissue) was around 25-30% shorter in *H. defensa*-infected aphids compared with uninfected aphids (Table 1; Fig. 1A). The total number of intracellular punctures (pd) made by *H. defensa* infected aphids was around two-fold higher than those made by uninfected aphids (Table 1; Fig 1B). Furthermore, following the first stylet probe into plant tissue, the first intracellular puncture (pd) occurred more rapidly for infected aphids (Table 1). Although the frequency of intracellular punctures increased in *H. defensa* infected aphids (Table 1; Fig. 1B), the duration of these intracellular punctures was on average 50% shorter for infected aphids compared with uninfected aphids (Table 1; Fig. 1C). The frequency of these intracellular punctures was highest and their duration shortest in the first hour (Table 1). Following this, the frequency of intracellular punctures in the second to sixth hours was not affected by endosymbiont presence, although the duration of intracellular punctures was influenced by symbiont presence in the second and sixth hours of EPG monitoring (Table 1).

Three aphid feeding parameters relating to stylet activity in the vascular tissue were affected by endosymbiont presence (Table 1; Fig. 1D-F): aphids infected with *H. defensa* showed a 50% reduction in time spent salivating into the phloem during stylet contact with the phloem (Table 1; Fig. 1D), displayed a 33% increase in phloem ingestion during stylet contact with the phloem (Table 1; Fig. 1E) and had a higher proportion of phloem sap ingestion phases (E2 phases) containing a period of sustained phloem sap ingestion (sE2 - a period of ingestion > 10 mins) (Table 1; Fig. 1F).

### Interactive effects of plant partial-resistance against aphids with aphid endosymbiont infection

In line with previous findings (Leybourne et al., 2018), the mass gain of *R. padi* nymphs was reduced when feeding on the partially-resistant wild relative of barley Hsp5 compared with aphids feeding on the susceptible modern cultivar of barley Concerto (ANOVA plant species: F_1,93_ =122.57; p = <0.001; Fig. S1), although endosymbiont presence/absence alone did not affect aphid fitness (ANOVA endosymbiont: F_1,93_ = 0.42; p = 0.514). The growth of aphids infected with *H. defensa* was further reduced by 22% when aphids were feeding on Hsp5 (ANOVA plant species x endosymbiont interaction: F_1,93_ =6.35; p = 0.013; Fig. S1). To examine whether alterations in aphid probing and feeding behaviour contributed towards this fitness cost, we identified EPG parameters responding differentially to endosymbiont infection on each plant type.

Fifteen EPG parameters were significantly affected by the endosymbiont infection x plant type interaction (Table 2). Eleven of these were differentially affected by *H. defensa*-infection for aphids contained on Hsp5 (Table 2). These data indicated that, in support of our second hypothesis (that differential aphid probing and feeding behaviour between uninfected and *H. defensa*-infected aphids is a key contributor towards the decreased fitness of *H. defensa*-infected aphids feeding on partially-resistant Hsp5), altered aphid probing and feeding behaviour could contribute towards decreased fitness of *H. defensa*-infected aphids on this less suitable plant (Fig. S1). When interacting with Hsp5, infected aphids spent 9% less time probing into plant tissue compared with uninfected aphids (Table 2), resulting in an overall reduction in the total time spent probing into plant tissue (Table 2; Fig. 3A). Although there was no difference in the number of non-probing phases between *H. defensa*-infected and uninfected aphids when feeding on Hsp5 (Table 2), *H. defensa*-infected aphids spent longer periods not probing into the plant tissue (Table 2). Furthermore, the duration of the first probe into plant tissue by *H. defensa*-infected aphids feeding on Hsp5 was around six-fold shorter compared with uninfected aphids (Table 2; Fig. 2B), and *H. defensa*-infected aphids required twice as many probes into plant tissue before the phloem was reached compared with uninfected aphids (Table 2; Fig. 2C). The total time spent ingesting phloem was reduced by 44% for *H. defensa*-infected aphids feeding on Hsp5 compared with uninfected aphids (Table 2; Fig. 2D) and the longest observed period of phloem ingestion was three-fold shorter for *H. defensa*-infected aphids compared with uninfected aphids when feeding on Hsp5 (Table 2; Fig. 2E).

Infection with *H. defensa* also altered the feeding behaviour of *R. padi* when probing into the susceptible barley cv. Concerto: *H. defensa*-infected aphids achieved sustained phloem sap ingestion two-fold faster than uninfected aphids (Table 2; Fig. 2F), however this did not affect aphid growth (Fig. S1).

## Discussion

By analysing aphid feeding behaviour, our study provides novel mechanistic insights into the consequences of *Hamiltonella defensa* infection for interactions at the aphid-plant interface and shows that *H. defensa*-infection can alter aphid probing behaviour, irrespective of host plant suitability, with potential consequences for insect fitness. In addition to this, our data show that these interactions can be influenced by plant susceptibility to, or resistance against, aphids and we provide novel evidence showing that aphid physiological processes are differentially affected by endosymbiont presence and host plant suitability which, at least in part, explains a fitness cost associated with *H. defensa*-infection for *R. padi* when feeding on a poor quality (partially-resistant) host plant.

### Endosymbiont infection alters aphid exploratory probing into plant cells and promotes phloem ingestion

When probing into plant tissue, *H. defensa*-infected aphids displayed a characteristic pattern of more frequent and shorter exploratory intracellular punctures (EPG waveform pd) than uninfected aphids. The precise cause of this symbiont-associated effect on aphid probing is not clear, although a similar pattern was recently reported in *H. defensa*-infected cowpea aphids, *Aphis craccivora* (Angelella et al., 2018), and it is likely that changes in intracellular puncture frequency will affect the transmission of plant viruses (Fereres and Collar, 2001; Powell, 2005). A key difference between our study and the previous work of Angelella et al. (2018) was that *R. padi* infected with *H. defensa* also showed differential feeding behaviour caused by altered stylet activities within the phloem. *H. defensa*-infected aphids spent less time salivating into the phloem and showed an overall increase in the percentage of phloem phases which contained phloem ingestion, including an increased proportion of these containing periods of sustained phloem ingestion. Altered aphid probing and feeding behaviour did not appear to affect aphid fitness directly as no overall effect of *H. defensa* infection on *R padi* growth, development, fecundity, or longevity was detected (present study; Leybourne et al., 2018). However, *H. defensa*-infection can affect aphid fitness in other species (Zytynska et al., 2019 preprint) and differential feeding behaviour in *H. defensa*-infected aphids could be associated with these altered aphid phenotypes, including an increased adult body mass and enhanced offspring production in black bean aphids, *A. fabae* (Castañeda et al., 2010). Endoymbiont-induced changes in feeding behaviour might be due to indirect effects of the bacterium on stylet activities mediated by bacterium-derived salivary factors (Su et al., 2015; Frago et al., 2017).

The extent of these endosymbiont-derived fitness consequences can often be dependent on aphid clonal line or aphid genotype (Castañeda et al., 2010) and it is important to note that endosymbiont-conferred traits vary between different aphid lines, aphid genotypes, and aphid species (Castañeda et al., 2010; Vorburger and Gouskov, 2011; Leybourne et al., 2018): indeed, *H. defensa*-infection can also reduce *A. fabae* reproductive rate and survivorship (Vorburger and Gouskov, 2011) and decrease *S*. *avenae* fecundity (Li et al., 2018). Altered probing behaviour might also explain differential plant responses to infestation by aphids infected with *H. defensa*, including changes in the emission of Herbivore Induced Plant Volatile (HIPV) compounds (Frago et al., 2017) and reduced dry matter allocation to roots (Hackett et al., 2013; Bennett et al., 2016). A focus for future research should include the consequences of aphid species and genotype for *H. defensa*-associated modifications to aphid probing and feeding behaviour to fully elucidate their effects on aphid pest status, virus transmission, and plant responses to aphid infestation.

### Endosymbiont infection reduces aphid feeding on a poor quality host plant

When probing into the partially-resistant plant, Hsp5, aphids infected with *H. defensa* showed a differential physiological feeding pattern compared with uninfected aphids, including a reduction in the time spent probing into plant tissue, an increase in the number of plant tissue probes required to reach the phloem tissue, and a decrease in total phloem ingestion. This was linked with decreased fitness in *H. defensa*-infected aphids compared with uninfected aphids when feeding on Hsp5, in line with our previous findings (Leybourne et al., 2018). A decrease in the duration of the first probe into plant tissue, and an overall reduction in time spent probing into the plant tissue, are representative of mesophyll- and epidermal-derived factors which inhibit and impede the penetration of the aphid stylet through the plant tissue, as highlighted by Alvarez et al. (2006). A similar fitness cost associated with *H. defensa*-infected aphids has been observed previously in *A. fabae* feeding on different quality plant species (Chandler et al., 2008), although it is not known if this was linked with altered aphid probing and feeding behaviour.

We recently characterised the partial-resistance mechanism of Hsp5 (Leybourne et al., 2019) and reported that partial-resistance involves mesophyll and phloem traits. These included an increased abundance of defensive thionins and a reduction in the availability of essential amino acids as mesophyll-derived and phloem-derived partial-resistance factors, respectively (Leybourne et al., 2019). These factors could underlie the decreased time aphids spent probing into plant tissue and the shorter duration of the initial probe into plant tissue, although the underlying processes causing these differential feeding patterns are currently unclear. A key factor which likely contributes towards this decrease in aphid fitness is our observation that *H. defensa*-infected aphids showed a 44% reduction in time spent ingesting phloem on Hsp5 compared with uninfected aphids. It is probable that this substantial decrease in phloem ingestion contributes significantly to the 22% reduction in nymph growth we detected. Indeed, a previous study using the peach-potato aphid, *Myzus persicae*, have shown that a 58% decrease in ingestion rate can result in a 10% reduction in aphid growth (Karley et al., 2002). We also detected differential feeding patterns between *H. defensa*-infected aphids feeding on Hsp5 and Concerto: *H. defensa*-infected aphids feeding on Concerto achieved sustained phloem feeding more rapidly than those feeding on Hsp5. A faster initiation of sustained feeding could explain the higher mass of *H. defensa*-infected nymphs on Concerto. However, it is likely that our observed reduction in nymph mass for both infected and uninfected nymphs when feeding on Hsp5 compared with those feeding on Concerto is due increased aphid resistance in Hsp5 (Leybourne et al., 2019). The rapid initiation of sustained feeding could be associated with other aphid fitness effects which are currently uncharacterised, such as the transmission or acquisition of phloem-limited viruses.

### Conclusion

In this study, two hypotheses were tested: 1) that infection with *H. defensa* can lead to altered aphid probing and feeding behaviour; 2) that differential aphid probing and feeding behaviour between uninfected and *H. defensa*-infected aphids is a key contributor towards the decreased fitness of *H. defensa*-infected aphids feeding on partially-resistant Hsp5. *R. padi* infected with the defensive facultative endosymbiont, *H. defensa*, showed altered probing and feeding behaviour compared with uninfected aphids, irrespective of plant type, including an increase in the number of intracellular punctures and in phloem ingestion, supporting our first hypothesis. Furthermore, in support of our second hypothesis, our EPG data highlight novel mechanistic processes which contribute towards an observed fitness cost arising from *H. defensa*-infection in *R. padi* feeding on the partially-resistant plant, Hsp5, which was associated with a reduction in the time aphids probe into the plant tissue, an increase in the number of plant tissue probes required to reach the phloem, and a 44% reduction in total phloem ingestion. Together, our results show that aphid facultative endosymbionts can influence aphid-plant interactions in more subtle ways than previously realised and indicate that plant suitability can exacerbate these effects.

## Acknowledgements

We would like to thank A. Nicholas E. Birch (James Hutton Institute) for helpful comments on the manuscript.

## Author Contributions

AJK and DJL conceived and designed the experiments. DJL performed the experiments and analysed the data. All authors contributed to data interpretation. DJL wrote the manuscript with input from all authors. All authors read and approved the final manuscript.

